# Novel elasticity measurement techniques reveal that the *C. elegans* cuticle is required for physical integrity with age

**DOI:** 10.1101/2021.10.04.463135

**Authors:** Mohammad Rahimi, Salman Sohrabi, Coleen T. Murphy

## Abstract

Changes in biomechanical properties have profound impacts on human health. *C. elegans* might serve as a model for studying the molecular genetics of mammalian tissue decline. Previously, we found that collagens are required for insulin signaling mutants’ long lifespan and that overexpression of specific collagens extends wild-type lifespan. However, whether these effects on lifespan are due to mechanical changes during aging has not yet been established. Here, we have developed two novel methods to study the cuticle: we measure mechanical properties of live animals using osmotic shock (OS), and we directly perform the tensile test (TT) on isolated cuticles using microfluidic technology. Using these tools, we find that cuticle, not the muscle, is responsible for changes in ‘stretchiness’ of *C. elegans*, and that cuticle stiffness is highly non-linear and anisotropic. We also found that collagen mutations alter integrity of the cuticle by significantly altering elasticity. Additionally, aging stiffens the cuticle under mechanical loads beyond the cuticle’s healthy stretched state. Measurements of elasticity showed that long-lived *daf-2* mutants were considerably better at preventing progressive mechanical changes with age. These tests of *C. elegans* biophysical properties suggest that the cuticle is responsible for their resilience.

## Introduction

The study of cell and tissue mechanics is crucial in understanding physiological and pathological processes and how they change with age. Mechanical forces play a major role in stem cell differentiation^1^, cell migration^2^, and cellular polarization^3^. Moreover, most pathological conditions, such as orthopedic disorders, atherosclerosis^4^, cancer^5^, and neurodegenerative disorders^6^ are associated with significant changes in the mechanical characteristics of cells. Aging, in particular, is associated with mechanical changes in tissues that may reduce overall life quality or accelerate aging^7, 8^. For instance, skin loses its elasticity, bones become brittle, and muscles lose mass. Stiffening of the vasculature with age is associated with cardiovascular disease, as well.

*C. elegans* is a premier model organism for studying age-related changes because of its high genetic tractability, optically transparent body, relatively simple cellular structure, and short life span. Like higher-order organisms, *C. elegans’* epithelial cells are essential for development and regulation of nutrients and ions coming from the outside environment. Hypodermal cells are the main component of the epithelial system and secrete the cuticle, which wraps around the whole body and forms an extremely flexible and resilient extracellular matrix (ECM). The nematode’s cuticle is mainly composed of cross-linked collagens and additional insoluble proteins that are synthesized in hypodermal cells^9^. Collagen provides strength, elasticity, and structural stability to fibrous tissues. Mutations in individual collagen genes result in defects in worm morphology (e.g., Dumpy and Blister mutants) and can cause embryonic and larval death^10^. We previously discovered that the expression of many cuticle collagens that decrease with age remain high in long-lived *daf-*2 insulin signaling mutants and are required for its longevity. Moreover, the overexpression of some of these collagen genes also extends the life span of wild-type animals^11^. However, the mechanism by which these collagens alter the worm’s lifespan are not clear.

To dissect molecular mechanisms of changes in worms’ mechanical properties, we need a robust technique to measure the cuticle’s stiffness. Various techniques have been developed to probe the mechanical properties of cells and tissues, including piezos^12^, force probes^13^, micropipette aspiration^14^, optical tweezers^15^, atomic force microscopy (AFM)^16^, magnetic twisting cytometry^17^, and microelectromechanical systems (MEMS)-based force clamp^18^. Among them, a pressurized microfluidic chip^19^ and AFM^20^ have been used to probe *C. elegans* cuticle mechanics. Using volumetric compression measurements by applying hydrostatic pressure, it was suggested that the bulk mechanical properties of whole worms are independent of the cuticle, and that collagen mutants targeting the cuticle do not impact the bulk modulus^19^. In contrast, a recent AFM study of *C. elegans* stiffness and topography suggested that there is a gradual loss in cuticle stiffness with age, and dietary restriction regimes directly influence the biomechanical properties of animals^20^. However, AFM measurements are highly localized and can have considerably different elastic moduli^20^, and may result in irreproducibility, depending on the probe and contact mechanics model used. Thus, a more reliable technique for measuring the mechanical properties of worms may provide a better understanding of the role of the cuticle stiffness changes during *C. elegans* aging.

Here, we developed osmotic shock and tensile test techniques to measure stress-strain relationship and elasticity of the cuticle at the whole-body and isolated cuticle scales, and upon disruption by collagen mutation. We then investigated how mechanical strength correlates with age and in longevity mutants. Our results suggest that the cuticle’s elasticity is highly nonlinear and anisotropic. Also, the elasticity of the *C. elegans* cuticle changes considerably with age, and this change is robustly delayed in the long-lived *daf-2* insulin receptor mutant.

## Results

### Shrinking and swelling during osmotic shock

The cuticle is under hydrostatic pressure from internal tissues^19^. The cuticle’s anisotropic mechanical properties and the worm’s internal pressure determine its shape in homeostasis (Figure 1a). The worm’s cuticle is semi-permeable to water, and upon transferring to the hypo-osmotic solution, the worm swells until it explodes (Supplementary Video 1). We can also reduce the internal pressure by applying a hyper-osmotic shock as the water exits and the body shrinks in an anisotropic manner (Supplementary Video 2). A time-course analysis of the changes in body shape was used to estimate the cuticle’s stiffness based on these changes in shape.

**Figure 1.**
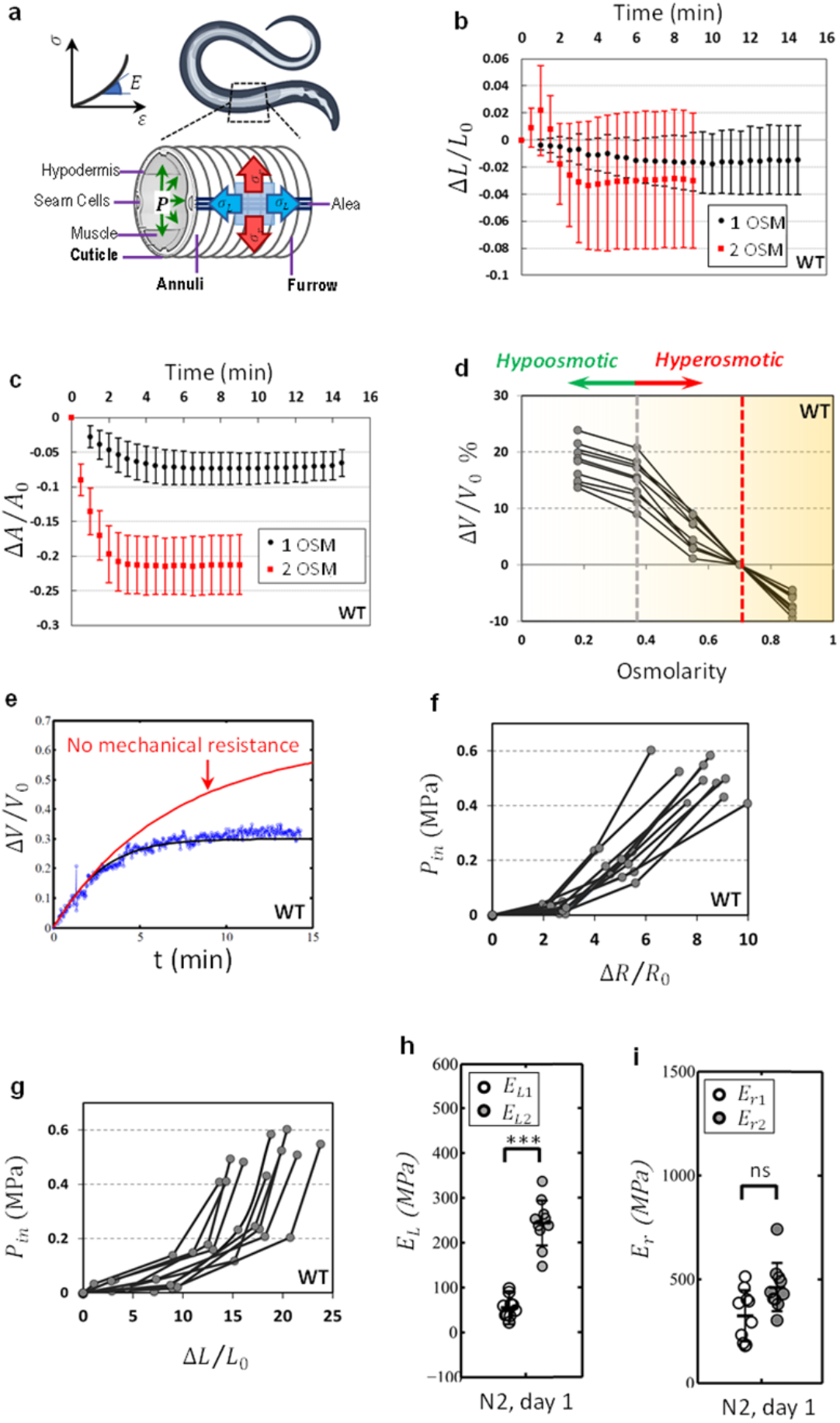
Anisotropic and nonlinear elasticity of *C. elegans*. (a) Internal pressure stretches *C. elegans*’ cuticular structures (Annuli, Furrow, and alae). (b) and (c) Time-course measurement of worms’ length and projected area after transfer to high osmolarity solutions. Higher osmolarity solutions generate more overall shrinkage and it may take up to 20 min for water exchange to reach the steady state. *n*=10 per condition. (d) Wild-type animals are transferred from hyperosmotic media to hypo-osmotic media quasi-statically. *V*_0_ is the volume of the flaccid worms. *n*=10 per condition. (e) Time-course measurement of volume upon transferring worms to lower osmolarity solution (0.55 OSM to 0.25 OSM). Applying a mathematical model to these data points, we can estimate the mechanical force resisting swelling. *V*_0_ is the initial volume of the animal before the osmotic shock. (f) and (g) Using our mathematical model and body shape change data, the internal pressure can be calculated. *n*=10 per condition. Elastic moduli in radial (h) and longitudinal (j) directions for wild type worms under small and large stretching loads. *n*=10 per condition. Two-tailed t-tests. ***p <0.*001*. Plots show 25th percentile, median, and 75th percentile.

The internal and external osmolarities reach equilibrium by exchanging water with the outside environment through its semi-permeable cuticle. Volume exchange can be described as

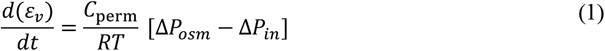

where *R* is the Boltzmann constant, *T* is temperature, *C*_*perm*_ is the permeability, Δ*P*_*osm*_ is osmotic pressure, and *P*_*in*_ is internal pressure. Osmotic pressure can also be rewritten as

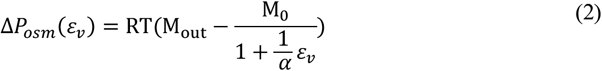

where M_0_ is internal osmolarity, M_out_ is the osmolarity of the environment, and *α* is the water fraction of the worm’s volume. *M*_out_ = 0.345 *OSM* is the healthy concentration for the worm’s environment either in fluid or in agarose gel^21^. Since the cuticle becomes wrinkled and flaccid at high osmolarities, the internal pressure of worms at 1 *OSM* and 2 *OSM* solutions are assumed to be zero. We can record the projected area and length of worms to quantify body shape changes as animals are transferred to these high osmolarity solutions (Figure 1b-c). From there, normalized volume changes can be calculated using *ε*_*v*_ = Δ*V*/*V*_0_ = (Δ*A*/*A*_0_) + (Δ*R*/*R*_0_) to estimate *α* = 0.65 and M_0_ ≅ 0.7 *OSM* using Equation 2. We then measured volume changes of animals with respect to their flaccid state at different osmolarities upon gradually transferring them from hyperosmotic media to hypo-osmotic buffer (Figure 1d). In our mathematical model, if there is no mechanical resistance (Equation 1), the animal would exchange water until it reaches solution osmolality (Figure 1e). The difference between this prediction and our real-time volume change measurement demonstrates how much mechanical load is imposed on the cuticle in the form of internal pressure (Figure 1f-g).

### Anisotropic non-linear behavior of the cuticle

Our calculations of internal pressure demonstrated that it only slightly changes as *M*_out_ is increased to 0.55 *OSM* (Figure 1f-g). The additional increase in internal pressure resulted in non-linear responses in length and radial changes under small and large stretching loads. The cuticle is always under this small stretching load when maintained on growth media with healthy osmolarity. The larger stretching load happens during hypo-osmotic shock, where the cuticle is stretched beyond its normal stretched state. By simplifying the cuticle as a thin cylinder which is the load-bearing component for internal pressure (Supplementary Figure 1a), we can plot stress-strain curves and accordingly calculate the elasticity of the cuticle at different stretching loads (Supplementary Figure 1b-c). We found that the cuticle is generally stiffer in the radial direction compared to the longitudinal direction, consistent with the presence of most structural collagen fibers, which are oriented in the azimuthal direction (Figure 1h-i). Moreover, due to a regular pattern of annular furrows on the cuticle in the longitudinal direction, the cuticle behavior is highly nonlinear under small and large stretching loads (Figure 1h). By contrast, the radial elasticity was not significantly different between these stretching loads (Figure 1i).

The cuticle’s response to hypo-osmotic shock can be discussed in terms of stiffness quantifiable by elastic moduli (*E*_*L*2_ and *E*_*r*2_) (Figure 2-3). We also demonstrated cuticle response to hyper-osmotic shock using body shape changes (Δ*L*/*L*_0_ and Δ*R*/*R*_0_) that were described in term of resistance. To complement this, we also showed overall elasticity of cuticle under small stretching load using bulk moduli, κ_1_ (Figures 2f, 3f).

**Figure 2.**
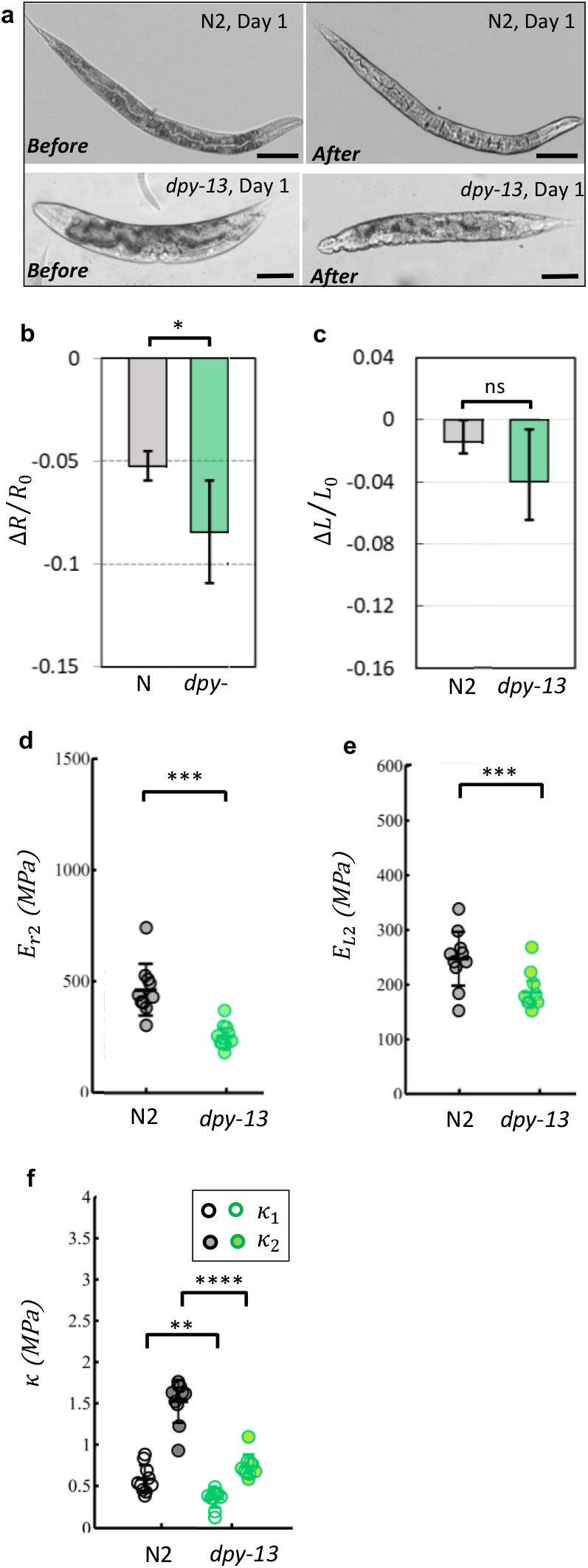
Collagen mutants have a softer cuticle. (a) Young wild-type and *dpy-13* worms’ immediate response to hyperosmotic shock (0.35 OSM to 1 OMS). The appearance of wrinkles in the young collagen gene mutant indicates a lower cuticular resistance. Scale bar = 100 μm. (b) and (c) Worms are transferred to 1 OSM solution and imaged after 20 min. Length and area changes of wild-type and *dpy-13* worms at Day 1 are calculated; N2/day 1: *n=*12, *dpy-13*/day 1: *n*=16. (d) and (e) Elastic moduli in radial and longitudinal directions for young wild type and collagen mutants. N2/day 1: *n=*10, *dpy-13*/day 1: *n*=10. (f) Bulk moduli of wild-type and collagen mutants under small and large stretching loads. The cuticle defective mutant is significantly softer than wild type. N2/day 1: *n=*10, *dpy-13*/day 1: *n*=10. Two-tailed t-tests. *p < 0.05, **p < 0.01, ***p < 0.001, ****p < 0.0001. Plots show 25th percentile, median, and 75th percentile.

**Figure 3.**
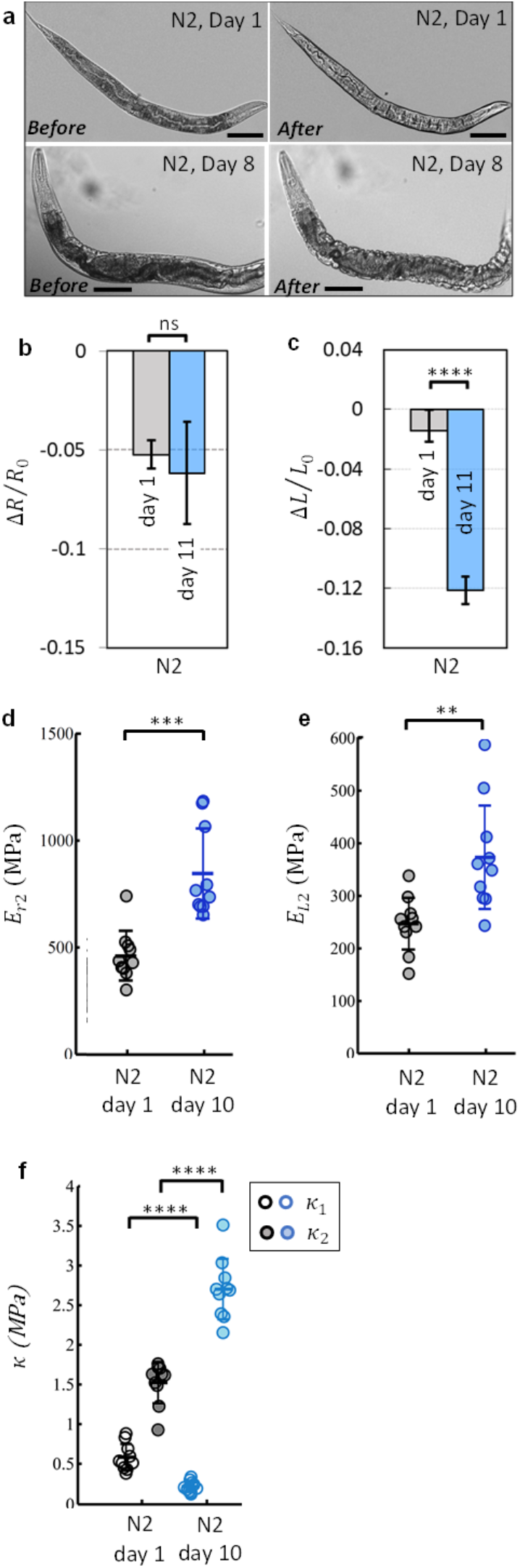
Aging alters the stiffness of the *C. elegans* cuticle. (a) Young and aged wild-type worms’ immediate response to hyperosmotic shock (0.35 OSM to 1 OSM). Scale bar = 100 μm. (b) and (c) Worms are transferred to 1 OSM solution and imaged after 20 min. Length and area changes of wild-type worms at Day 1 and Day 11 are calculated; N2/day 1: *n=*12, N2/day 11: *n*=8. (d) and (e) Elastic moduli in radial and longitudinal directions for young and aged wild type worms; N2/day 1: *n=*10, N2/day 10: *n*=10. (f) Bulk moduli of young and aged wild-type worms. The cuticle softens at small stretching loads and stiffens at larger stretching loads. N2/day 1: *n=*10, N2/day 10: *n*=10. Two-tailed t-tests. *p < 0.05, **p < 0.01, ***p < 0.001, ****p < 0.0001. Plots show 25th percentile, median, and 75th percentile.

### Collagen mutations alter integrity of the cuticle

To investigate how collagen contributes to the elasticity and strength of cuticle, we have tested collagen mutants. Mutations in cuticle collagen genes with homozygous reduced or loss-of-function alleles, including *dpy-13*, result in a phenotype described as Dumpy (Dpy) and have narrower annuli than those of the wild-type worms^22^. Upon transferring young adult animals from M9 buffer to high osmolarity solution, we found that the cuticle of young *dpy-13* mutants immediately became wrinkled compared to wild-type cuticle (Figure 2a). The larger radial shrinkage (Figure 2b) and significantly more elastic bulk modulus, κ_1_ (Figure 2f) of young *dpy-13* mutants indicated a lower hyperosmotic resistance compared to wild-type animals (Figure 2). Similarly, under hypo-osmotic swelling, *dpy-13* also showed significantly lower radial and longitudinal stiffnesses compared to wild type (Figure 2d-e). These findings indicated that the lack of some level of collagen expression leads to a softer cuticle (Figure 2f), which may explain some of the mutant’s defects.

### Osmotic shock analysis of aged cuticles

Aging has a significant effect on the anisotropic mechanical properties of the cuticle (Figure 3a). We observed that aged (Day 11) wild-type worms shrink considerably in the longitudinal direction in response to hyper-osmotic shock (Figure 3c). Similarly, bulk moduli under small stretching loads also indicated softening of cuticle with age (Figure 3f). By contrast, our measurements of high stretching load with hypoosmotic swelling showed that the cuticle becomes significantly stiffer in both directions with age (Figure 3d-e and Figure 3f).

### Tensile test of isolated cuticles

Measurements of whole worms or intact worm bodies suggest that changes in the cuticle are responsible for the worm’s stiffness, but do not rule out contributions from muscles or other tissues. Thus, we needed to directly measure the stiffness of the worm’s cuticle to confirm that the osmotic shock technique measures changes in the cuticle. To do so, we first isolated intact cuticles from the worms^28^. We built a 3-layer microfluidic chip to stretch isolated cuticles, and measured the cuticle’s membrane deflection *h* as a function of the applied pressure (Figure 4a-b). The push-up valve is first pressurized to measure the deflection of membrane without cuticle before initiating the tensile test (TT) (Figure 4c). Then, the isolated cuticle, free of cellular and membranous material, was adhered to the thin PDMS membrane and push-down valves were pressurized to clamp the cuticle (Supplementary Figure 1d). The drop in the membrane deflection with respect to deflection without specimen presented the cuticle resistance against stretching (Figure 4c).

**Figure 4.**
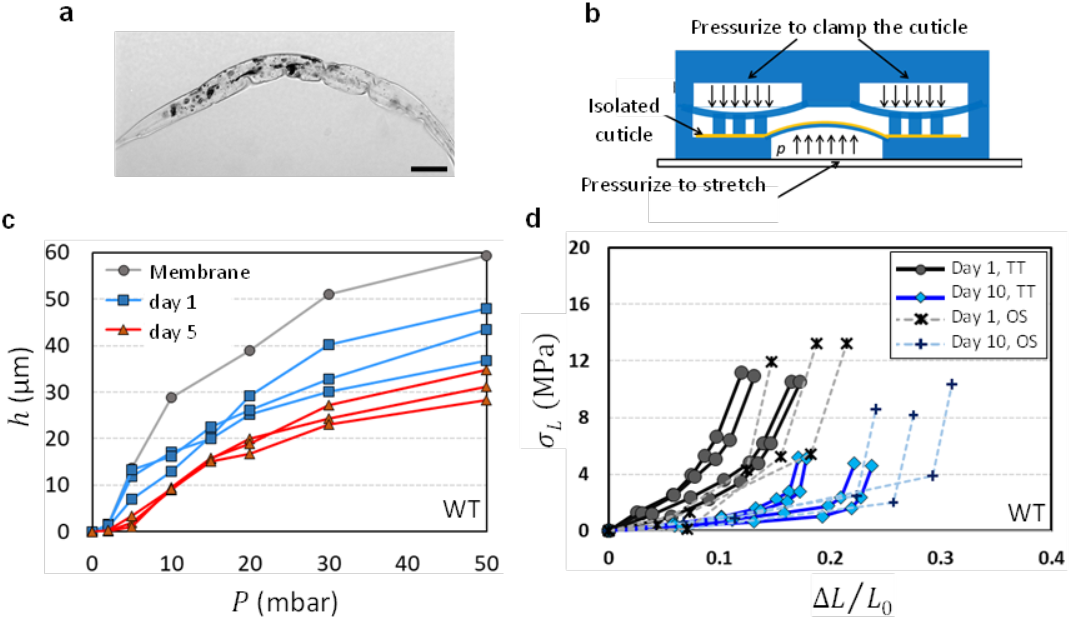
Tensile test of the cuticle using a microfluidic device. (a) The isolated cuticle is free of cellular and membranous material. Scale bar = 100 μm. (b) Side view of a 3-layer chip to apply stretching force on an isolated cuticle. (c) The PDMS membrane was deflated using a pressure pump with and without the cuticle. The membrane deflection *h* is recorded as a function of the applied pressure P. Day 1 wild type has a more flexible cuticle than Day 5 worms; N2/day 1: *n=*3, N2/day 5: *n=*3. (d) Marginal agreement of stress-strain curves acquired using the tensile test with osmotic shock measurements. Aged (Day-10) wild-type worms have higher stiffness than Day-1 worms. N2/day 1/TT: *n=*4, N2/day 10/TT: *n=*4, N2/day 1/OS: *n=*3, N2/day 10/OS: *n=*3.

### Tensile test measurements of young and aged cuticle

Compared to the Day 1 isolated cuticle, the isolated cuticle of Day 5 worms exhibited more resistance against membrane deflection. Thus, these measurements suggest that Day 1 wild-type animals have a more flexible cuticle in comparison with a Day 5 wild-type animal’s cuticle, indicating that cuticle stiffness increases with age. Our osmotic shock measurements in the longitudinal direction can be compared with the results from the direct tensile test of the isolated cuticle by calculating normalized length change using Δ*L*/*L*_0_ = (*θ*. *R* – *a*)/*a*, where *R* = (*h*^2^ + *a*^2^)/2*h*, and sin(*θ*) = *a*/*R* (Supplementary figure 1e). We found that stress-strain curves of isolated cuticles corroborate the results of our osmotic shock results (Figure 4d). The slope of the stress-strain curve indicates that both techniques reflect a stiffening of the cuticle with age at larger stretching loads.

### Osmotic shock analysis of long-lived *daf-2* mutants

We next tested long-lived *daf-2* insulin receptor mutants to determine whether their longevity is correlated with the mechanical characteristics of the cuticle. *daf-2* mutants increase lifespan through their regulation of a suite of genes that affect proteostasis, innate immunity, stress responses, fat stores, and metabolism, among other processes^23–27^. We observed that aged longevity mutants were considerably better at maintaining resistance under hyperosmotic shock at youthful levels (Figure 5a and Figure 5b-c). Similarly, stiffness changes with age under swelling in both directions were robustly slowed in *daf-2,* while the wild-type cuticle becomes considerably stiffer (less elastic) with age (Figure 5d-e).

**Figure 5.**
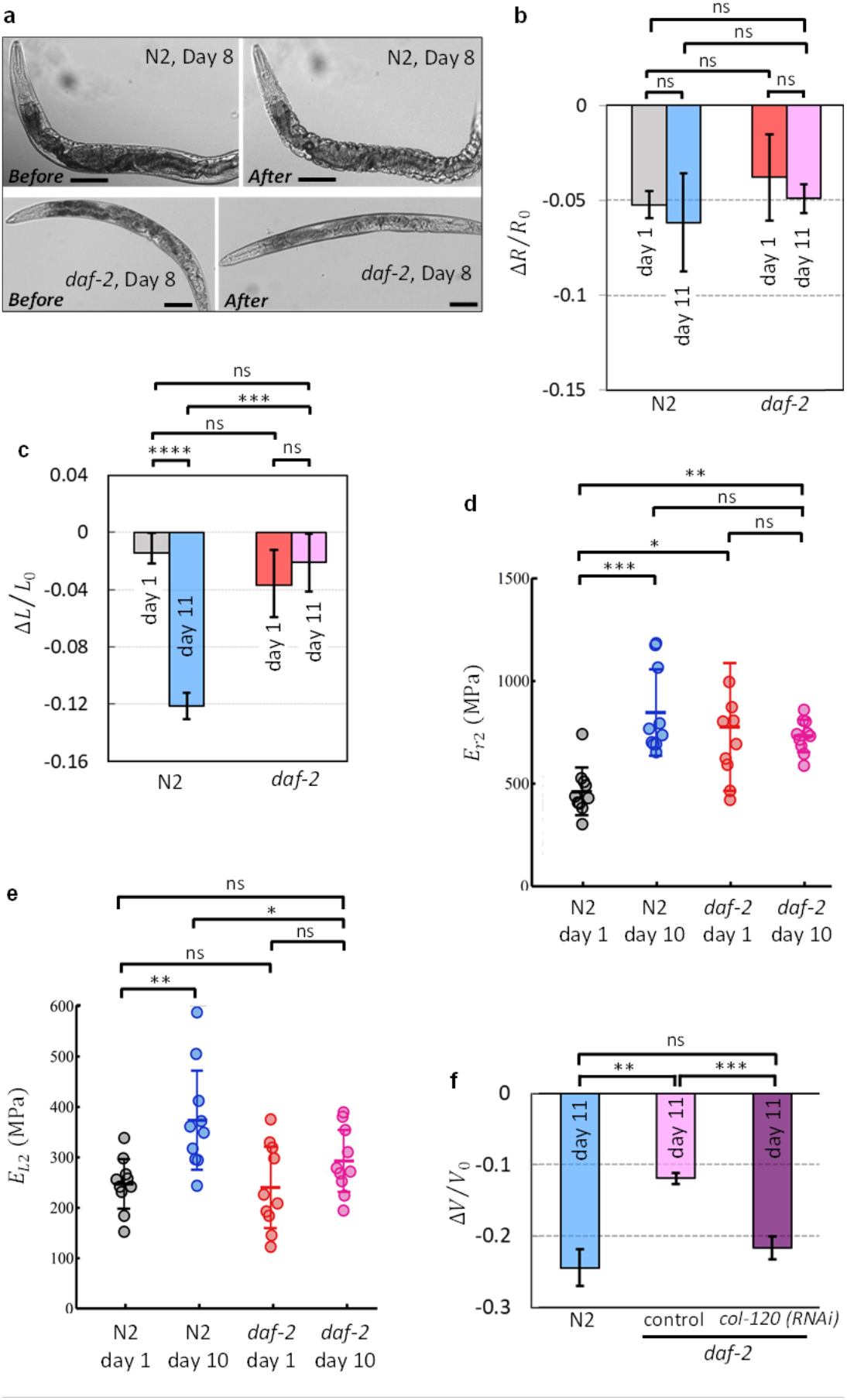
Stiffness changes with age is slowed in the *daf-2* longevity mutant. (a) Aged wild-type and *daf-2* worms’ immediate response to hyperosmotic shock (0.35 OSM to 1 OSM). Absence of wrinkles in the aged long-lived mutant signals better cuticle resistance. Scale bar = 100 μm. (b) and (c) Worms are transferred to 1 OSM solution and imaged after 20 min. Length and area changes of wild-type and *daf-2* worms at Day 1 and Day 11 are calculated. Decreasing resistance with age is slowed in *daf-2* mutant; N2/day 1: *n=*12, N2/day 11: *n*=8, *daf-2*/day 1: *n=*12, *daf-2*/day 11: *n*=8. (d) and (e) Elastic moduli in radial and longitudinal directions for young and aged wild type and long-lived mutants. Increasing stiffness of cuticle with age is slowed down in *daf-2* mutant. N2/day 1: *n=*10, N2/day 10: *n*=10, *daf-2*/day 1: *n=*10, *daf-2*/day 10: *n*=10. (f) Adulthood collagen gene knockdown reduced *daf-2* ability to resist volumetric shrinkage upon transferring to 1 OSM solution. *n*=8 for N2 and *daf-2 (control)*. *n*=14 for *daf2*(*col-120 RNAi*). One-way ANOVA with Dunnett’s post-hoc. *p < 0.05, **p < 0.01, ***p < 0.001, ****p < 0.0001. Plots show 25th percentile, median, and 75th percentile.

We previously showed that reduction of collagen gene expression during adulthood reduced *daf-2* lifespan, and the overexpression of collagens extends wild-type lifespan11; therefore, we tested whether the reduction of collagens that are required for *daf-2’s* long lifespan alters the mechanics of the epithelial system. RNA interference knockdown of *col-120* in the *daf-2* mutant decreased *daf-2’s* resistance to volume loss to wild-type levels in aged (Day 11) worms (Figure 5f), suggesting that collagens are critical for the extended maintenance of elasticity in *daf-2* mutants.

## Discussion

Changes in mechanical properties with age can have profound impacts on health. For example, human arteries, which are scaffolded by elastin and collagen, progressively increase in stiffness by 200–250% between the ages of 25 and 75 years^29^, and this increased stiffness is associated with the development of cardiovascular disease. Aging causes collagen matrix to unravel, total collagen content to increase, cross‐ linking to occur, and collagen fiber distribution to disorganize, all of which lead to dysfunction, including arterial stiffening^30, 31^.

*C. elegans* is well-suited for the study of age-related changes in tissue mechanics^11^. However, few methods existed for studying cuticle mechanics. Here, we have introduced two novel, complementary techniques that allowed us to measure the mechanical properties of the cuticle. Osmotic shock is a high-throughput technique for whole-worm measurement of stiffness with robust reproducibility (Supplementary video 3). By quantifying the body shape change and calculating the worm’s internal pressure, a simple mathematical model can be used to estimate the anisotropic mechanical properties of the cuticle. To confirm our results from osmotic shock, we developed a more direct method to measure stiffness using microfluidic technology to stretch an isolated cuticle, to measure the cuticle’s longitudinal stiffness.

Previous measurements of cuticular local stiffness using a piezoelectric transducer indicated a high elastic modulus in the 380 MPa range^12^. Another similar local stiffness study utilizing the buckling of nanowires reported an elastic modulus of 457 MPa on the lateral alae, and 257 MPa in the region immediately around the lateral alae^32^. Our measurements of wild-type animals averaged at *E*_*L*2_ ≅ 250*Mpa* and *E*_*r*2_ ≅ 450*Mpa*, which is in close agreement with these studies.

It was suggested that the bulk mechanical properties of whole, paraformaldehyde-fixed worms are independent of the cuticle^19^. However, the excessive muscular stiffness induced by high paralytic concentrations or by fixation might have obscured these measurements, as we found the cuticle rather than the muscle is the main component determining the stiffness of the worm. We also showed that despite its relative thinness (roughly 2% of the diameter of the worm), the cuticle is responsible for whole-worm changes in ‘stretchiness’ with age.

An AFM study of *C. elegans* stiffness and topography suggested that worms undergo a marked loss of stiffness and increased cuticle senescence during aging^20^. However, our results suggest that the cuticle’s behavior is highly non-linear; while the cuticle softens at small stretching loads similar to AFM measurements, it stiffens at large stretching loads. These findings were confirmed using both osmotic shock and direct tensile tests. It is possible that the spherical probe in the AFM setup is only able to measure cuticle behavior with highly localized small indentation forces and cannot deliver a broader understanding of the anisotropic nonlinear mechanical properties of the whole worm or whole cuticle^20^. These AFM measurements also showed that reduction of the IGF-1 signaling pathway preserved stiffness with age^20^. Similarly, we also found that long-lived *daf-2* mutant worms were better at preventing progressive softening and stiffening with age under small and large stretching loads, respectively. Therefore, we believe that the approaches we describe here have provided a more comprehensive understanding of cuticle mechanics. We also tested a *Dumpy* mutant to investigate how defects in microstructure affect the mechanics of whole tissue and found the cuticle of *dpy-13* worms is significantly softer than wild-type cuticles. Additionally, knocking down the extracellular structural genes that preserve the life span of longevity mutants altered the physical properties of the *C. elegans* exoskeleton and reduced the resistance of *daf-2* animals in response to osmotic shock.

Detailed mechanical studies of *C. elegans* will enable the characterization of a largely unexplored phenotype that has important consequences for human aging. Specifically, cuticle stiffness can be used as a biomarker for healthspan. Based on their specific attributes, these assays can be used to assess changes in body frailty of worms *in vivo* to probe the role of specific collagens in the regulation of longevity, pathogenesis, and osmotic stress regulation. Our results suggest that cuticle components that help animals retain their “youthful” mechanical properties, particularly elasticity, with age. This elasticity may not only be a result of maintained youthfulness, but may also contribute to longevity. Overall, this work will provide the platform to integrate molecular genetics with biophysical measurements, providing a more complete understanding of the frailty of worms as they age.

## Materials and Methods

### *C. elegans* strains and maintenance

*C. elegans* strains were grown at 20°C on nematode growth medium (NGM) plates or high growth medium (HG) plates seeded with OP50 *Escherichia coli*. RNAi clones were obtained from the Ahringer RNAi library. The following strains were used in this study: wild-type worms of the N2 Bristol strain, *daf-2(e1370),* and *dpy-13(e184).*

### Osmotic shock

Worms were immobilized by 100 μM sodium azide in M9 buffer. Then they were deposited in high osmolarity solution by adjusting the concentration of sodium chloride. After 20 minutes, they were quasi-statically transferred from hyperosmotic media to hypo-osmotic buffer with 20 min pauses in each step to ensure water exchange with the outside environment through semi-permeable cuticle is complete.

### Cuticle Isolation

The cuticle isolation protocol described by Cox *et al.*^28^ was used. Briefly, washed nematodes were suspended in sonication buffer (10 mM Tris-HCI, pH 7 .4, 1 mM EDTA, 1 mM phenyl-methanesulfonyl fluoride [PMSF], and given ten 20-sec bursts of sonification. Cuticles were washed several times with sonication buffer and suspended in 1 ml of ST buffer (1% SDS, 0.125 M Tris-HCI, pH 6.8) and heated for 2 min at 100°C. Spun-down cuticles were extracted again with ST buffer as described.

### Microfluidic chip

The instructions for molding master fabrication, PDMS device fabrication, and measurement of closing pressures were adopted from Fordyce *et al.*^33^.

### Statistics

For all comparisons between two groups, an unpaired Student’s t-test was performed. For comparisons between multiple groups, One-Way ANOVA was performed with post-hoc testing. GraphPad Prism was used for all statistical analyses.

## Acknowledgements

We thank the *C. elegans* Genetics Center for strains (P40 OD010440) and the Murphy lab for discussion. This work was supported by the Glenn Foundation for Medical Research award to C.T.M. and a DP1 Pioneer (NIH) awards to C.T.M. C.T.M. is the Director of the Glenn Center for Aging Research at Princeton and an HHMI-Simons Faculty Scholar. M.R. and C.T.M. designed experiments. M.R. performed experiments, M.R., S.S., and C.T.M analyzed data. S.S. and C.T.M wrote the manuscript.

