## Supplementary information for "Novel elasticity measurement techniques reveal that the *C. elegans* cuticle is required for physical integrity with age"

**a**

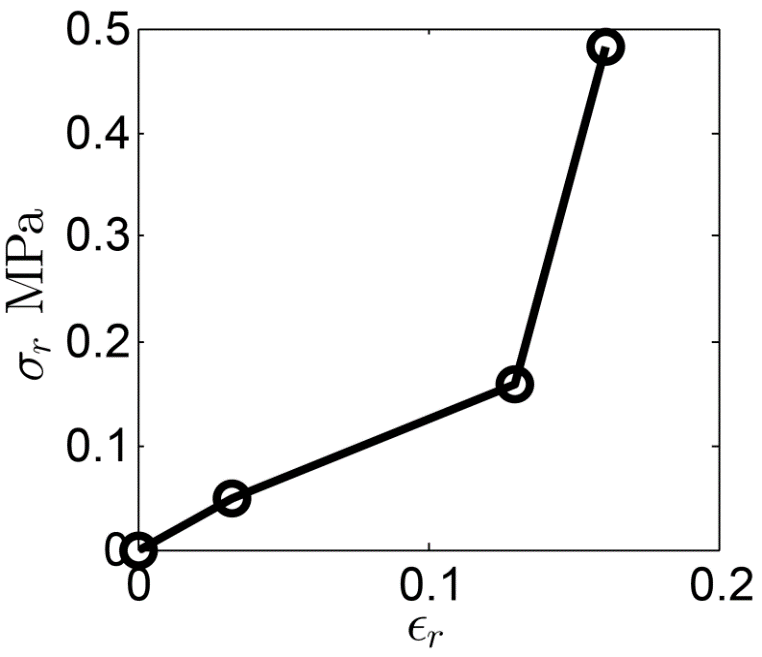

$$\boldsymbol{E}_{\boldsymbol{L}\boldsymbol{1}}$$

$$\boldsymbol{E}_{\boldsymbol{L}\boldsymbol{2}}$$

$\sigma_{L}$ (MPa)

$$\boldsymbol{\varepsilon}_{\boldsymbol{L}}$$

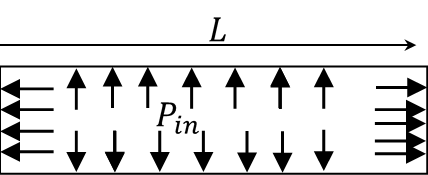

**b**

$\sigma_{r}$ (MPa)

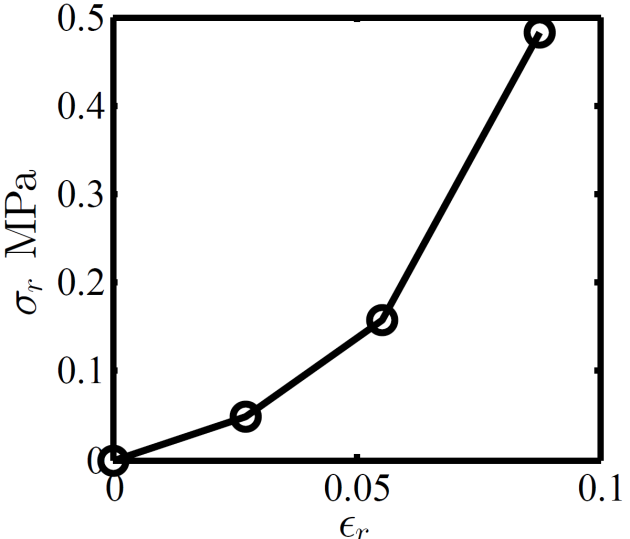

$$\boldsymbol{E}_{\boldsymbol{r}\boldsymbol{2}}$$

$$\boldsymbol{E}_{\boldsymbol{r}\boldsymbol{1}}$$

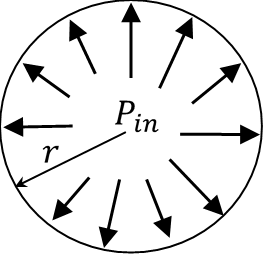

$$\boldsymbol{\varepsilon}_{\boldsymbol{r}}$$

**c**

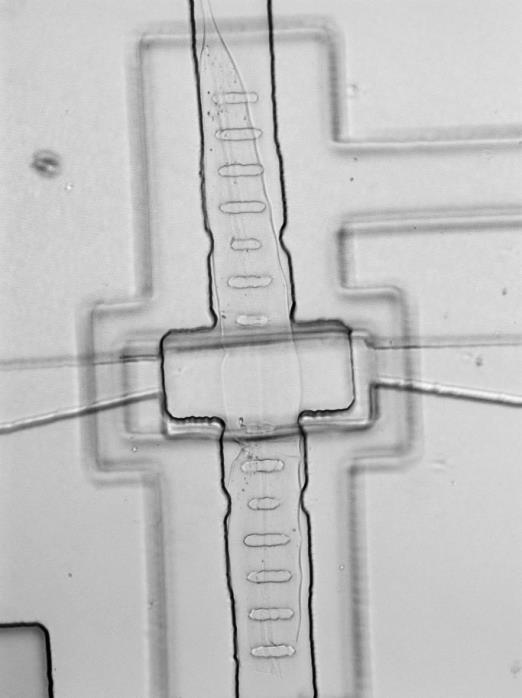

**d**

**Push down channel**

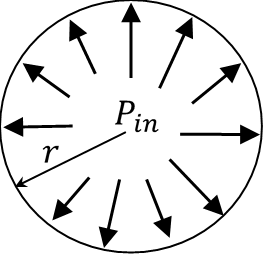

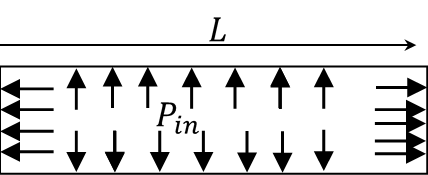

$$E_{r}=\frac{P_{in} rr_{0}}{t \Delta r}$$

$$E_{L}=\frac{P_{in} rL_{0}}{2 t \Delta L}$$

**e**

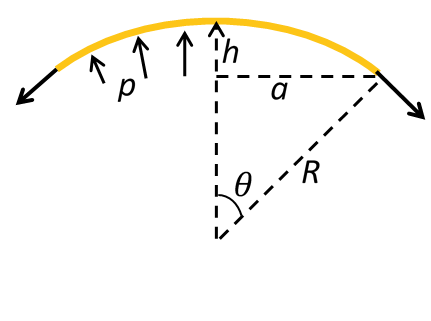

**Supplementary figure 1.** (a) A two-dimensional thin cylinder with anisotropic properties. Normalized radial can then be calculated as ${\Delta R}/{R_{0}}=\left( {\Delta A}/{A_{0}} \right)-\left( {\Delta L}/{L_{0}} \right)$. (b) and (c) Nonlinear elastic moduli of cuticle in longitudinal and radial directions under different loads can be derived from stress-strain curve. (d) The mechanism through which the inflation of thin membrane stretches isolated cuticle. (e) The schematics on how inflation heights at different applied pressure can be used to calculate the cuticle stiffness in longitudinal direction.

**Supplementary Video Legends**

**Supplementary video 1.** Hypoosmotic shock (0.35 OSM to DIW) make wild type (day 1) worm swell. The video is 10 times faster.

**Supplementary video 2.** Hyperosmotic shock (0.35 OSM to 1 OSM) make wild type (day 14) worm shrink. The video is 20 times faster.

**Supplementary video 3.** Osmotic shock as a simple and robust protocol for studying anisotropic mechanical properties of the whole worm can be carried out at high throughput in 96-well plates. Wild-type worms are transferred from 1 OSM to DIW. The video is 60 times faster.
